# A Cross-Sectional Study of *Staphylococcus Aureus* Colonization and the Nasal and Oropharyngeal Microbiomes

**DOI:** 10.1101/145540

**Authors:** Ashley E. Kates, Mark Dalman, James C. Torner, Tara C. Smith

## Abstract

**Background:** *Staphylococcus aureus* is a frequent cause of both infections globally. Colonization with the organism is known to increase the risk of developing infections and occurs in roughly one third of the general population. While many factors influence colonization, it has been demonstrated other members of the microbiome influence colonization with *S. aureus*. Here, we assessed the nasal and oropharyngeal microbiomes of healthy participants in relation to *S. aureus* colonization in a cross-sectional study using 16s rRNA sequencing of the v1-v3 region. As livestock workers have also been shown to be at an increased risk of carriage, we have also assessed microbiota differences in colonization status in a population of livestock workers.

**Results:** In both the nares and oropharynx, there were no microbiota differentially abundant between colonized and non-colonized persons. However, there was a significant difference in the beta diversity (Bray-Curtis distances) between carriers and non-carriers (*P*=0.002). When considering carriage stratified by livestock exposure, there were a number of differences. Most notably, colonized livestock workers had significantly more *Porphyomonas* (2-fold change = -8.54, *P* = 0.03) than the non-colonized livestock workers.

**Conclusions:** *S. aureus* is a frequent colonizer of the human upper respiratory tract, including the nares and oropharynx and causes a wide range of infections. Livestock workers are at increased risk for carriage. Interventions such as improving oral hygiene may lead to decreased *S. aureus* carriage by reducing other bacterial species such as *Porphyomonas*. Larger, longitudinal studies are needed to better explore what microorganisms may be associated with *S. aureus* colonization.

## Background

*Staphylococcus aureus* is an important pathogen globally and is traditionally associated with hospital-, community-, and most recently, livestock-associated infections. It is one of the most frequent causes of bacterial infections and has traditionally been the leading cause of skin and soft tissue infections [1]. It has been estimated roughly one third of the general population carry *S. aureus* [2] and 1.5% – 3% carry methicillin-resistant *S. aureus* (MRSA) [3]. While carriage itself is not necessarily harmful to the individual, it is a known risk factor for developing infections [4].

Colonization and infection with *S. aureus* has been extensively studied [5–7], predominantly in the hospital [8–10] and community [11, 12] settings; however, it remains unclear why some individuals persistently carry the bacteria while others do not, and why some individuals suffer repeated infections with *S. aureus* despite rounds of antibiotic treatments. Previous studies have shown other bacteria present on the body may decrease the likelihood of carrying *S. aureus* [13–16]. However, previous studies have been primarily based on traditional culture methods which underestimate the complexity of the microbiome.

In order to understand how microorganisms what bacteria may be important in preventing *S. aureus* colonization and infection, it is necessary to characterize what bacteria are present in the niches *S. aureus* inhabits. In this study we aim to determine what differences exist between carrier and non-carriers. Additionally, we aimed to determine if there were differences in the microbiota based on carrier status stratified by livestock exposure.

## Methods

### Study Population

Participants were enrolled into a cross-sectional study between April 2015 and March 2016 in Eastern Iowa. Participants were enrolled in their homes by the research team. A portion of the participants were enrolled from a prior, longitudinal study of *S. aureus* colonization. Participants enrolled from that study were selected to provide as even a number of colonized and non-colonized persons as possible. The remaining participants were enrolled from the community in one of 3 ways: through the Iowa Department of Natural Resources animal feeding operations database; a booth at Iowa county fairs; and lastly through snowball sampling where the research team asked already enrolled participants to invite community member they thought might be interested to participate. Eligibility criteria were: 18 years of age, speak English, have not taken antibiotics or inhaled corticosteroids in the prior three months, not had the nasal influenza vaccine in the last month, no active infections of the upper respiratory tract, no hospitalized for greater than 24 hours in the last three months, and did not have HIV/AIDS. We also requested participants not eat, drink, or brush their teeth within one hour of sample collection. The University of Iowa institutional review board approved all study protocols prior to recruitment.

After consenting, participants filled out questionnaires assessing demographic characteristics, medical history, and animal contact. Participants provided swabs from their anterior nares and oropharynx using sterile, dry, nylon flocked swabs (Copan Diagnostics, Murrieta, CA) collected by a trained researcher and transported to the University of Iowa Center for Emerging Infectious Diseases for processing. Bacterial DNA was isolated using the MO BIO PowerSoil DNA isolation kit (MO BIO Laboratories Inc, Carlsbad, CA) adapted for swab use by removing the swab head and placing it in the tube during bead beating. Negative controls (kit reagents only) were used for every batch of extractions. Samples were sent for sequencing (including library preparation) to the University of Minnesota Genomics Center. 16s rRNA sequencing of the v1-v3 region was done on the Illumina MiSeq using 2x300 nt reads.

### Statistical analysis

Sequences were assessed for quality using FastQC (Babraham Institute, Cambridge, UK) with poor quality reads filtered out (poor quality sequencing reads are defined as sequences with low base quality scores, short reads less than 200bp, reads with uncalled nucleotide bases, or any reads that could not assemble into paired reads). Reads were assembled using FLASh with the following parameters: minimum overlap = 30, maximum overlap = 150, and mismatch = 0.1 [17]. Adapters were removed from the merged file using Cutadapt [18]. USEARCH version 8.1.1861 and Python version 2.7.12 were used for chimera removal, operational taxonomic unit (OTU) binning, and taxonomy assignment at the genus level. The Ribosomal Database Project (RDP) classifier was used as the reference database. OTUs were grouped together based on 97% similarity. Any species level classification was done using BLAST+2.4.0 and the blastn function. Human-associated OTUs were also removed from the dataset using BLAST+2.4.0 and the blastn function.

The primary research question was to assess any difference in the microbiomes of *S. aureus* colonized persons compared to non-colonized persons in the nares and oropharynx, In addition to this, we assessed in any differences in the nasal and oropharyngeal microbiomes existed when also considering whether the subject had contact with livestock. This was done by stratifying the colonized and non-colonized persons by livestock contact as well as stratifying those with and without livestock contact by their colonization status.

R version 3.3.1 was used for all statistical analyses and plot generation using the following packages: phyloseq [19], vegan [20], DESeq2 [21], and ampvis [22]. Alpha diversity was assessed using the Inverse Simpson diversity index [23] and beta diversity was assessed using the Bray-Curtis dissimilarity measure [24]. Principal coordinates analysis (PCoA) was used to visualize beta diversity. Permutational multivariate analysis of variance (PERMANOVA), through the vegan package, was used to assess diversity differences between groups. PERMANOVA was chosen because it does not assume any distribution, unlike parametric tests [25]. The DESeq2 and ampvis packages were used to assess microbiota differences between groups. Results were considered significant if the *P* was less than 0.05.

## Results

### Participant demographics

Fifty-nine participants were enrolled into the study, 37 of which were colonized in either the nares, oropharynx, or both. Thirty participants were colonized in the nares and seven in the oropharynx. The average age of participants was 54.6 years (range: 28–85 years) and 69.5% were male. Fifty-eight participants identified as Caucasian (98.3%) and five identified as another race (8.5%). Twenty-six participants had contact with one or more types of livestock. Animal types included swine (n=18), cattle (n=12), poultry (n=4), sheep (n=4), horses (n=2) and goats (n=1).

### Microbiota comparisons by colonization status

There was no significant difference in inverse Simpson’s diversity index between colonized and non-colonized persons in either the nares (*P* = 0.376) or the oropharynx (*P* = 0.728) (Figure 1). The PCoA of the Bray-Curtis distances for all samples is shown in Figure 2. The samples cluster by both colonization status (*P* >0.001) and sample type (*P*>0.001). When considering only the nasal samples, the significant difference between the colonized and non-colonized persons remained (*P* = 0.002) (Figure 2b); however, there was no difference between the colonized and non-colonized clusters in the oropharyngeal samples (*P* = 0.899) (Figure 2c).

**Figure 1:**
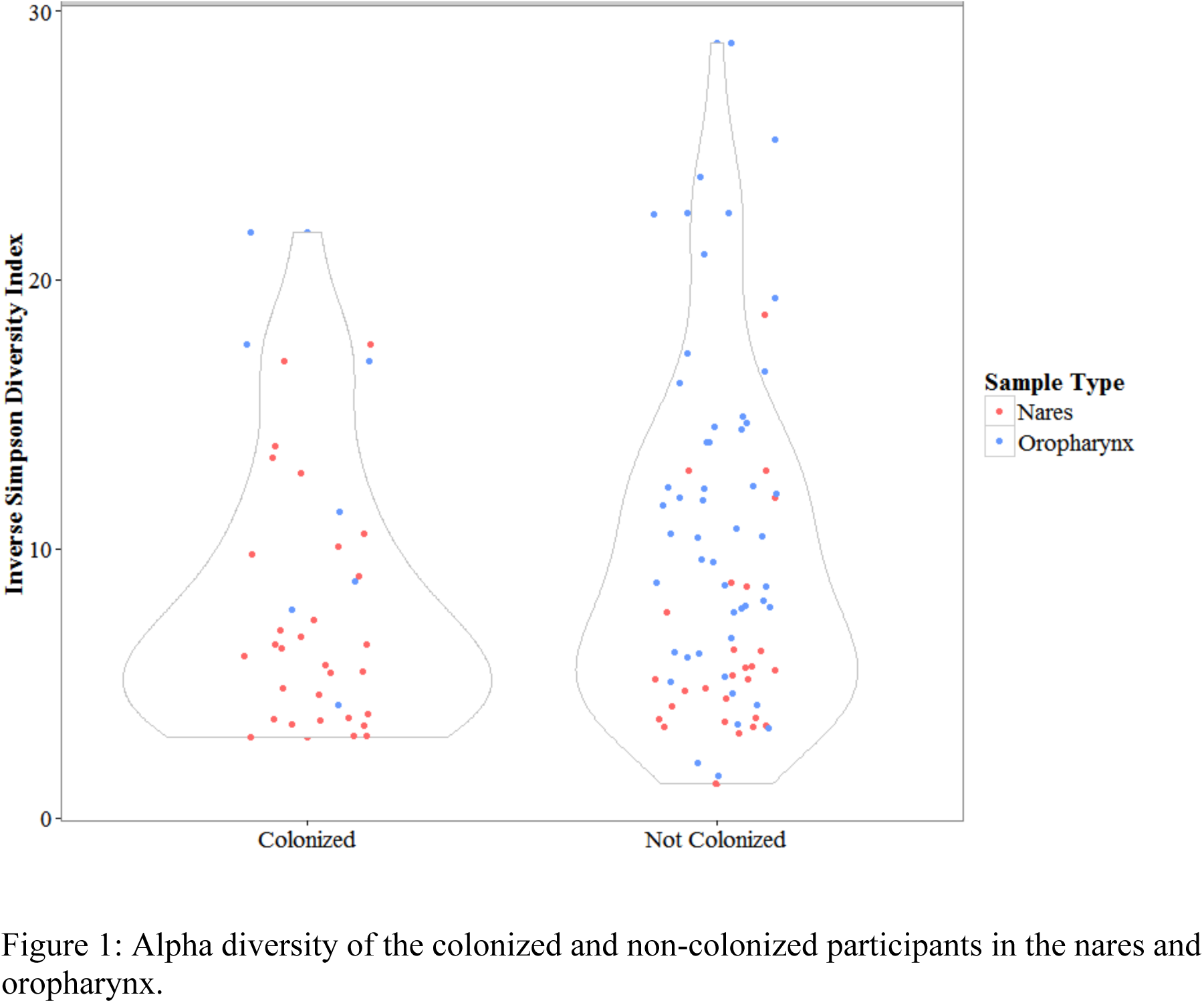
Alpha diversity of the colonized and non-colonized participants in the nares and oropharynx.

**Figure 2:**
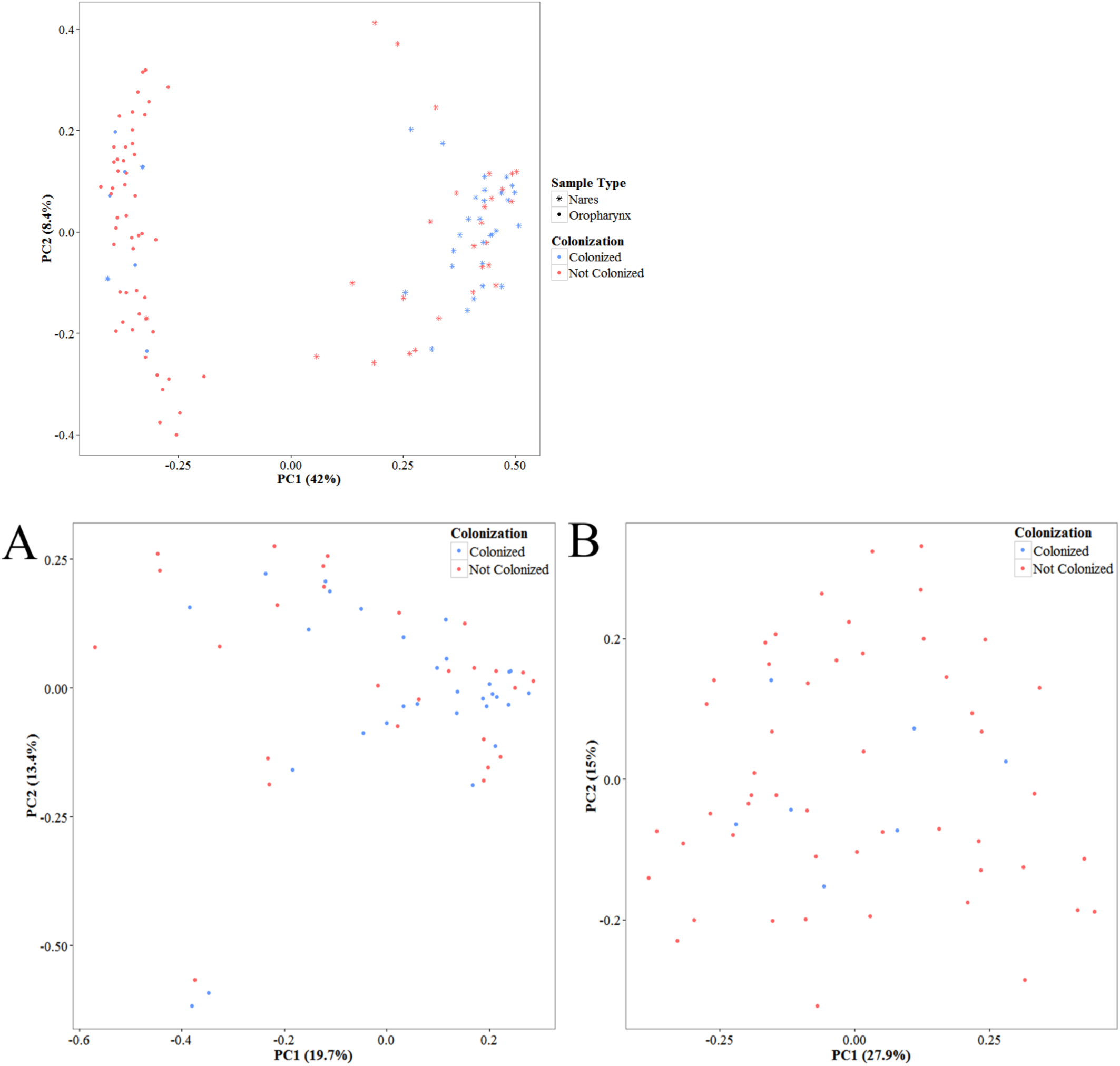
Principal coordinates analysis by colonization status. (a) Ordination based on the Bray-Curtis dissimilarities of the nasal and oropharyngeal sample microbiomes. (b) Ordination based on the Bray-Curtis dissimilarities of the nasal sample microbiomes. (c) Ordination based on the Bray-Curtis dissimilarities of the oropharyngeal sample microbiomes. PC1 and PC2 = principal coordinates 1 and 2, respectively.

Figure 3 shows the relative abundances of all OTUs in the colonized and non-colonized persons in the nares and oropharynx. There is a great deal of similarity between colonized and non-colonized person’s microbiota in the nasal samples, though there are several OTUs that are abundant in a greater number of colonized samples. The oropharyngeal microbiomes of colonized persons and non-colonized persons look very similar. OTUs belonging to the Firmicutes and Actinobacteria phyla dominate the microbiomes of the colonized and non-colonized persons; however, colonized persons have a greater amount of OTUs belonging to the Firmicutes phylum.

**Figure 3:**
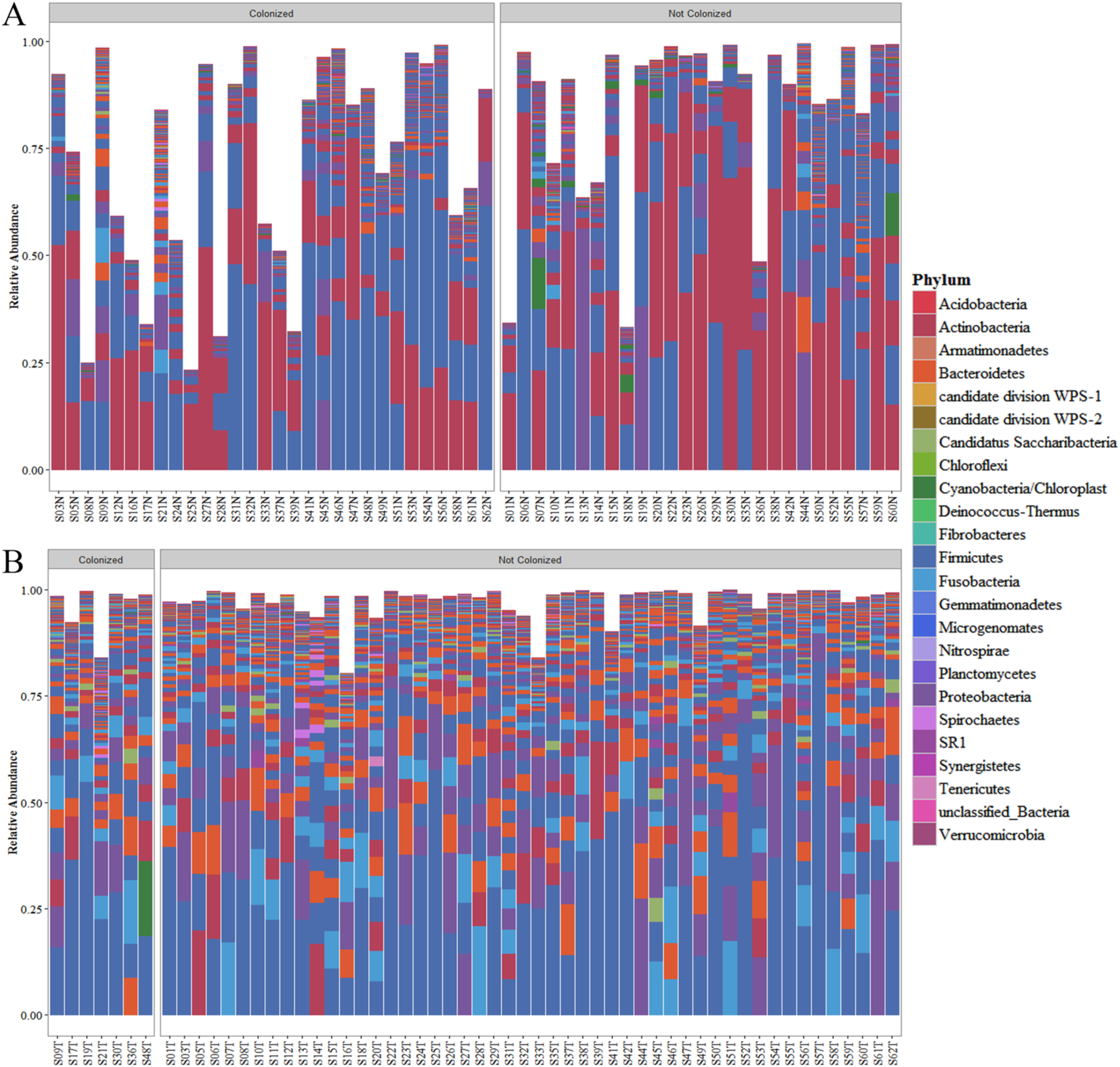
Relative abundances of bacterial phyla between colonized and non-colonized participants in the (a) nares and (b) oropharynx.

In the nasal microbiome, there was only one differential OTU, *S. aureus*. While the other *Staphylococcus* species were not significantly more abundant in the colonized persons, colonized persons did have more *Staphylococcus* OTUs compared to those not colonized with *S. aureus*. Other *Staphylococcus* species present were *S. epidermidis, S. cohnii, S. lentus, S. hominis, S. lugdunensis,* and *S. massiliensis*. *S. epidermidis* was the most prevalent non-*aureus* S*taphylococcus* species and was present in 100% (n=30) of the *S. aureus* colonized nasal samples compared to 89.6% (n=26) in the non-colonized samples. *S. cohnii* was the second most prevalent at 76.7% (n=23) of *S. aureus* colonized persons and 62.1% (n=18) of non-colonized samples. *S. homini* was prevalent in 60% (n=18) of colonized persons and 55.2% (n=16) of non-colonized persons. The largest difference between colonized and non-colonized persons was with *S. lentus* at 20% (n=6) in the colonized compared to 6.9% (n=2) in the non-colonized. Colonized persons also had slightly higher amounts of Proteobacteria (*Haemophilus* and an unclassified genus), Actinobacteria (*Rothia*), and Bacteroidetes (*Prevotella*). Non-colonized persons had higher amounts of Actinobacteria (*Corynebacterium*) and Cyanobacteria/Chloroplast (*Streptophyta*) compared to the colonized (Figure 4a).

**Figure 4:**
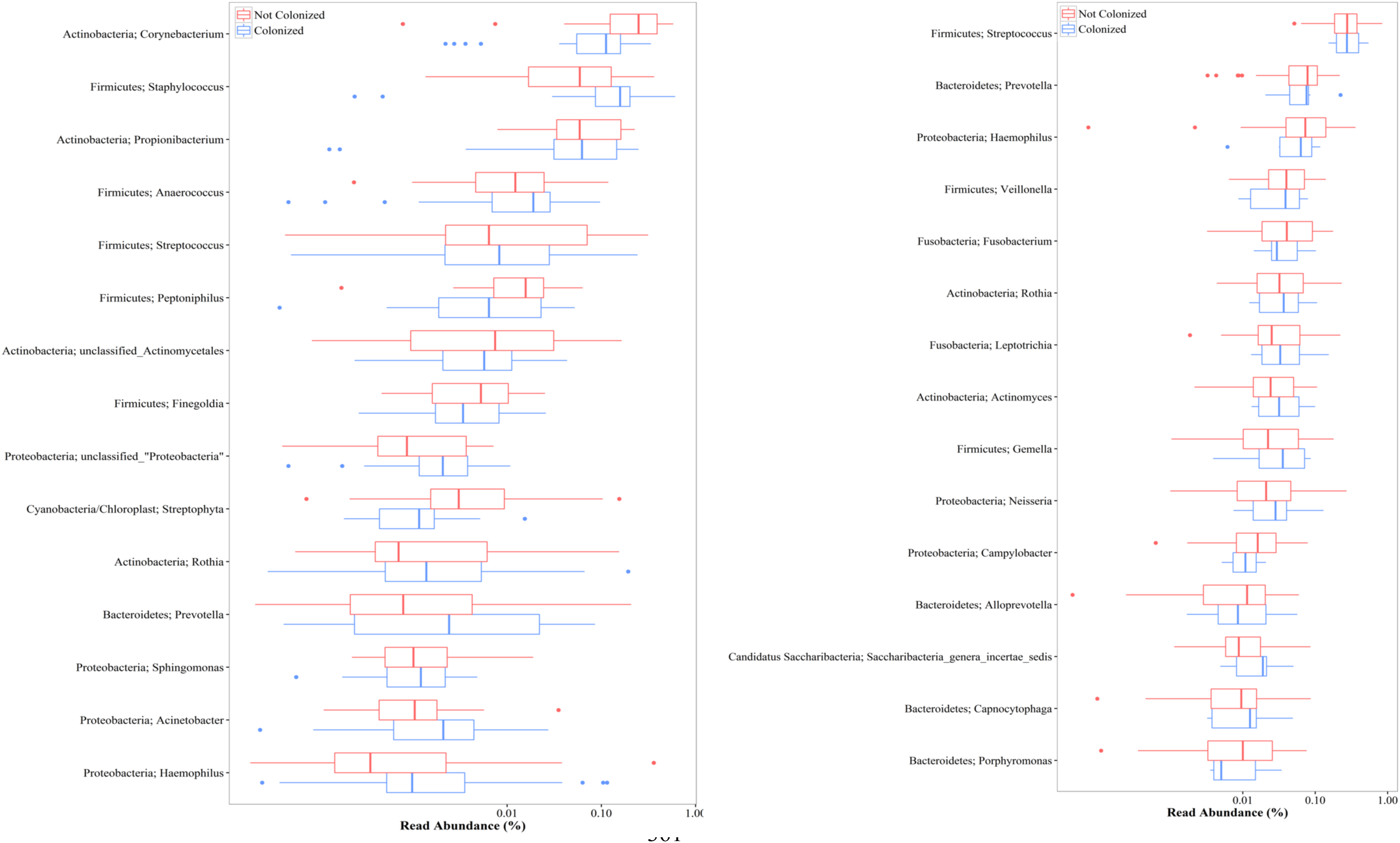
Top 15 most abundant organisms in the (a) nares and (b) oropharynx.

There were no OTUs significantly different between the colonized and non-colonized persons in the oropharynx. Many of the most abundant OTUs were similarly abundant in both the colonized and non-colonized persons. *Saccharibacteria genera incertae sedis* was slightly more abundant in the oropharynx of colonized persons. Bacteroidetes (*Porphyomonas)* was slightly more abundant among non-colonized persons (Figure 4b). Of the *Staphylococcal* species, *S. epidermidis* was the most prevalent and was in 100% (n=7) in the oropharynx of colonized persons compared to 35.6% (n=21) of the non-colonized persons. *S. cohnii* was present in the oropharynx of 42.9% (n=3) of the colonized persons and 28.6% (n=2) of the non-colonized persons.

### Colonization stratified by livestock contact

The Inverse Simpson’s diversity index was significantly greater for livestock workers in the nares (*P* = 0.0015) though not different in the oropharynx (*P* = 0.571); however, there was no difference in livestock exposure based on colonization status in either the nares (*P* = 0.815) or the oropharynx (*P* = 0.484) (Figure 5a, b). The ordination plots of colonization status and livestock exposure in the nares show the samples significantly cluster by colonization status (*P* = 0.001) and livestock exposure (*P* = 0.001) (Figure 6a); however, there are no significant clusters in the oropharyngeal samples (Figure 6b). When considering only colonized persons, samples associated with livestock exposure clustered separately from those without livestock exposure (*P* = 0.004). Samples associated with livestock exposure cluster together in the non-colonized persons (*P* = 0.021) (Figure 6c, d).

**Figure 5:**
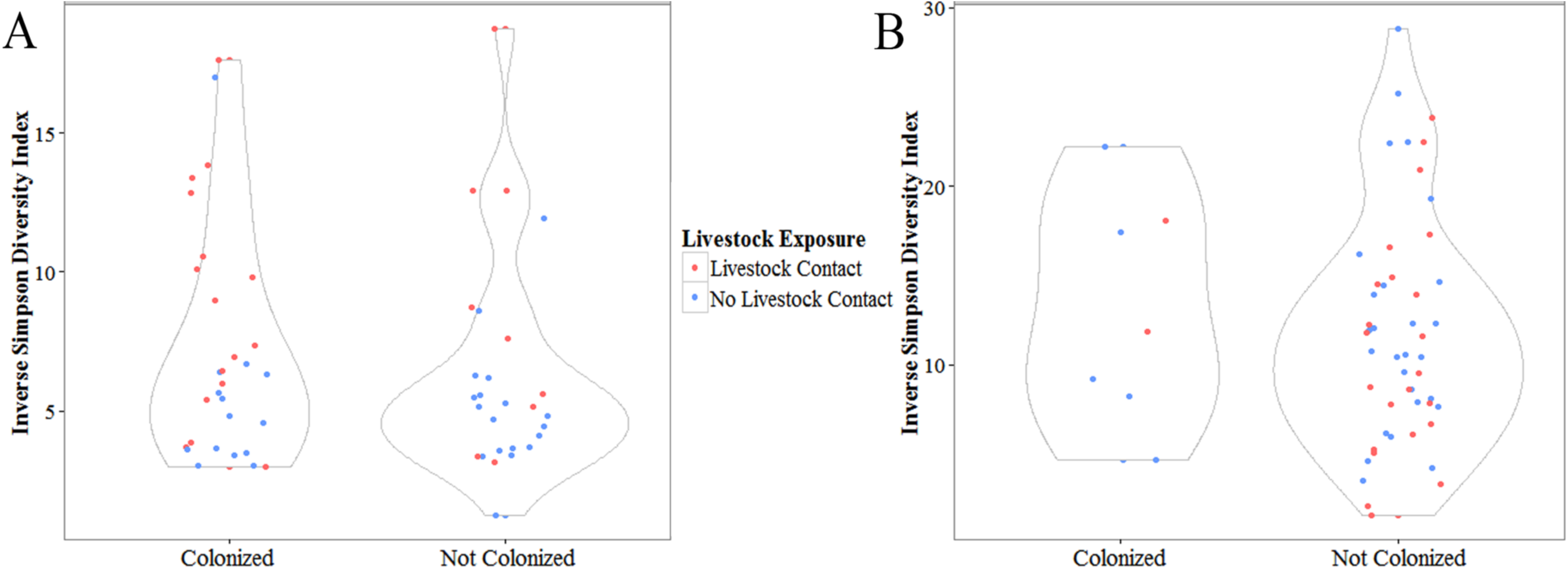
Inverse Simpson Diversity index by colonization status and livestock exposure in the (a) nares and (b) oropharynx.

**Figure 6:**
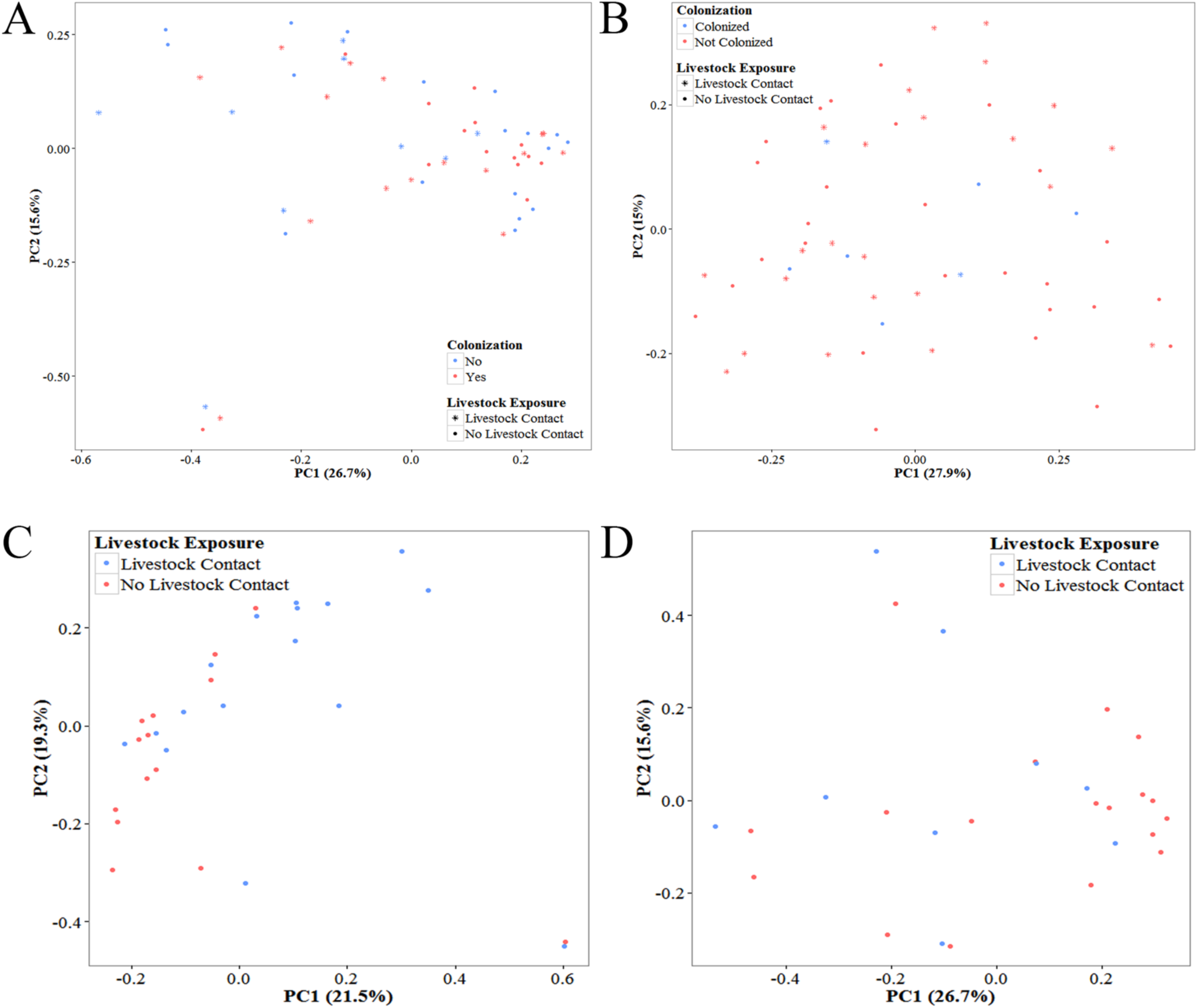
Principal coordinates analysis by colonization status and livestock exposure in the nares and oropharynx. (a) Ordination based on the Bray-Curtis dissimilarities of the all nasal samples. (b) Ordination based on the Bray-Curtis dissimilarities of all oropharyngeal samples. (c) Ordination based on the Bray-Curtis dissimilarities of the colonized nasal sample microbiomes. (d) Ordination based on the Bray-Curtis dissimilarities of the non-colonized nasal sample microbiomes. PC1 and PC2 = principal coordinates 1 and 2, respectively.

The most abundant OTUs in the nasally colonized persons belonged to the Firmicutes, Actinobacteria, and Proteobacteria. Fifteen OTUs were significantly differentially abundant between *S. aureus* colonized livestock workers and non-livestock workers. The majority of the significantly differential OTUs belonged to the Firmicutes phylum (Figure 7a). In the nasally non-colonized persons, Actinobacteria was the most abundant phylum followed by the Firmicutes, and Proteobacteria. Fewer differences were seen in abundance by livestock contact compared to the colonized persons. Seven OTUs were significantly more abundant in the non-colonized livestock workers compared to the non-colonized non-livestock workers and all belonged to the Firmicutes phylum (Figure 7b).

**Figure 7:**
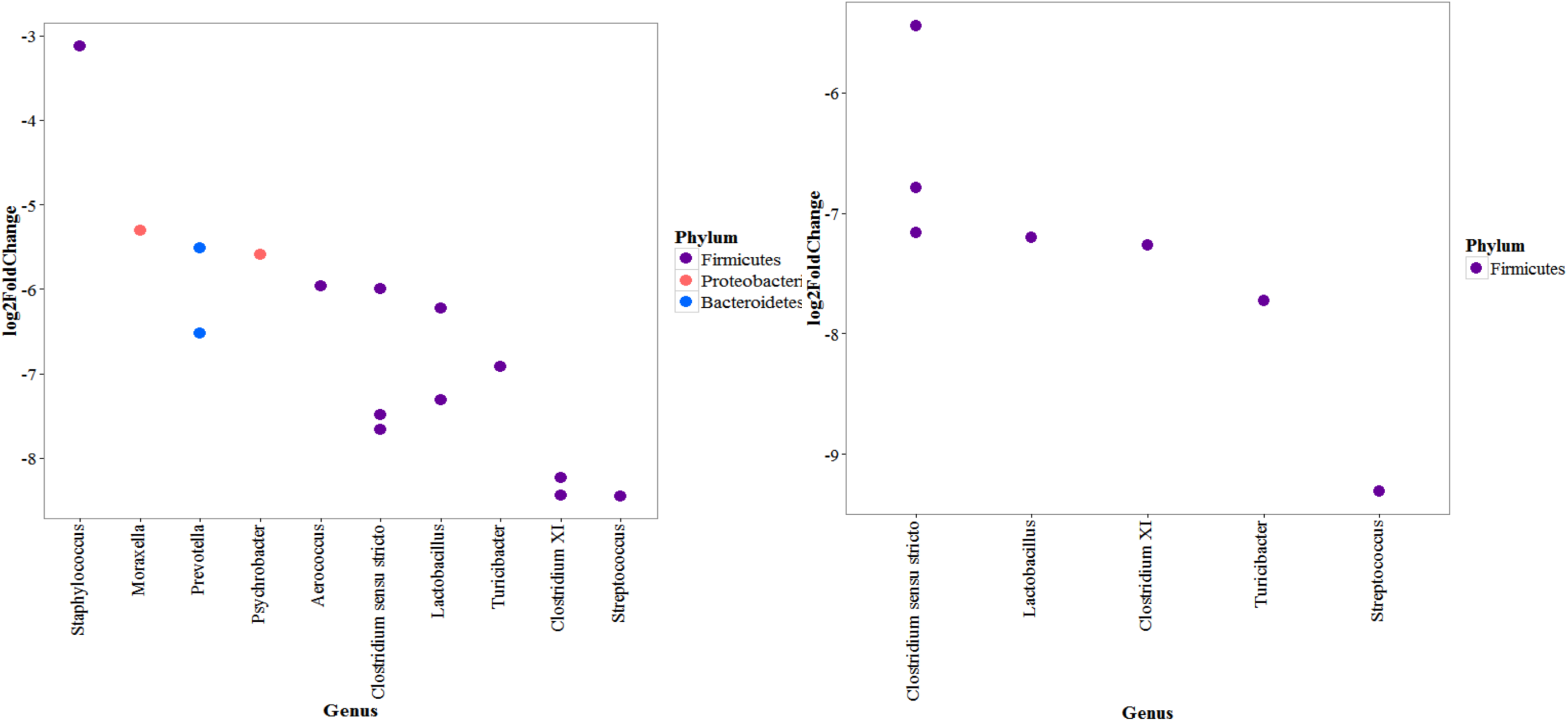
Differentially abundant organisms among colonized and non-colonized persons by livestock exposure. (a) Log 2-fold change of the significantly differentially abundant OTUs between colonized livestock workers and colonized persons with no livestock contact in the nares (Benjamini-Hochberg correction applied). Points represent OTUs with phyla represented by color. Negative values represent OTUs significantly more abundant in colonized livestock workers and positive values represent OTUs significantly more abundant in colonized, non-livestock workers. (b) Log 2-fold change of the significantly differentially abundant OTUs between non-colonized livestock workers and non-colonized persons with no livestock contact (Benjamini-Hochberg correction applied). Points represent OTUs with phyla represented by color. Negative values represent OTUs significantly more abundant in non-colonized livestock workers and positive values represent OTUs significantly more abundant in non-colonized, non-livestock workers.

Livestock workers oropharyngeally colonized with *S. aureus* had less *Strepotoccus, Prevotella, Heamophilus, Campylobacter* and *Veillonella* than oropharyngeally colonized non-livestock workers and had greater abundancies of *Rothia, Fusobacterium, Neisseria,* and *Lachnoanerobaculum* than non-livestock workers. *Alloprevotella* was the only significantly differentially abundant OTU and was more abundant in oropharyngeally colonized, non-livestock workers (2-fold change: 5.17, adjusted *P* = 0.0006). Among the oropharyngeally non-colonized persons, the abundancies between livestock workers and those without livestock contact were very similar and there were not significantly differentially abundant OTUs.

When considering livestock exposure stratified by colonization status, there were no differences in OTUs between the colonized and non-colonized livestock workers with the exception of *S. aureus* in the nares. The same was true for those without livestock exposure in the nares. However, in the oropharynx, colonized livestock workers had significantly more *Porphyromonas* (2-fold change = -8.54, *P* = 0.03) than the non-colonized livestock workers. Additionally, non-colonized livestock workers had significantly more *Atopobium* (2-fold change = 6.25, *P*= 0.004) compared to colonized livestock workers. There were no differences in the microbiomes of colonized persons with no livestock contact compared to non-colonized persons without livestock contact.

## Discussion

Understanding the ecologic niche *S. aureus* lives in is an important step in understanding the biologic factors associated with colonization and infection prevention. Here we have compared the microbiomes of the anterior nares and oropharynx and their relation to *S. aureus* in 59 healthy Iowans. Our population was predominantly male. This was because an aim of this study was to assess livestock workers. In the US, the majority of livestock workers are older males [26]. Male sex has been associated with increased risk of *S. aureus* carriage in non-hospitalized patients [27]. *S. aureus* colonization occurred in the nares of 30 participants and the oropharynx of seven participants. While were unable to identify any OTUs significantly associated with either *S. aureus* presence or absence, there were several differences between carriers and non-carriers, particularly in the nasal microbiome.

*S. aureus* nasal carriers’ microbiomes were significantly different from the non-carriers with regard to beta diversity (*P* = 0.002), though no individual OTUs were identified as differential between the groups. However, colonized individuals have more Firmicutes compared to the non-colonized participants while the non-colonized participants have more Actinobacteria. This is similar to other studies which have found strong, significant inverse correlations between the presence of Firmicutes and Actinobacteria in the nares [28] and have found communities dominated by Actinobacteria – particularly *Propionibacterium* and *Corynebacterium* – have fewer staphylococci [29]. While the differences were not significant, we also found *Corynebacterium* to be more frequently present in non-carriers compared to carriers; however, *Propionibacterium* was very similar between the two groups. *Corynebacterium* is one of the most consistently negatively-associated genera with regard to *S. aureus* colonization [28–32]. *Corynebacterium* has been shown to inhibit the growth of *S. aureus* in culture, specifically with regard to *C. pseudodiphtheriticum* [30].

Several other studies have found *S. epidermidis* to be both positively [30, 33] and negatively associated with *S. aureus* colonization [29]. Here, we observed *S. epidermidis* to be present 100% of the time when *S. aureus* was present in both the nares and the oropharynx. *S. epidermidis* was the most prevalent *Staphylococcus* species present in 56 of the 59 nasal samples and 28 of the oropharyngeal samples. The study by Frank et al. that identified *S. epidermidis* as being negatively associated with *S. aureus* carriage (*P* = 0.004) was done on primarily patients in intensive care units (ICUs) (n= 42) and only five non-hospitalized participants. It has been shown ICU stays can change the composition of the microbiome [34] with large changes to the ratios of Bacteroidetes and Firmicutes being observed in the gut microbiome [35]. It is possible the ICU environment is affecting the relationship between *S. aureus* and *S. epidermidis* in the Frank et al. study. It has also been hypothesized *S. epidermidis* will only inhibit *S. aureus* colonization when *S. epidermidis* is able to express a serine protease, *esp* [36] and the positive association between *S. aureus* and *S. epidermidis* is only observed when the majority of the *S. epidermidis* strains present in the microbiome do not express *esp*. However, a recent study demonstrated while *esp* was ubiquitous in all *S. epidermidis* strains tested, there was no correlation with *S. aureus* inhibition [37]. Future studies are need comparing hospitalized patients to healthy persons to better understand the association between the two microorganisms.

In the oropharynx, we observed a great deal of genus richness which is consistent with other studies [28]. As with prior studies, we observed the oropharynx to be dominated by Firmicutes as well as a high prevalence of Bacteroidetes and Proteobacteria [38]. Our inability to see any differences in the oropharyngeal microbiomes between carriers and non-carriers is possibly due to the small number of oropharyngeally colonized individuals (n=7).

When comparing the colonized individuals, colonized livestock workers had twenty-four OTUs that were significantly more abundant than in the nasal microbiota of their colonized, non-livestock worker counterparts. Almost all of these OTUs belonged to the Firmicutes phylum and many have been found to be associated with livestock contact. It is possible that at least a portion of livestock workers identified as *S. aureus* carriers are not truly colonized, only contaminated, and that while they are continuously in the presence of aerosolized *S. aureus, S. aureus* is unable to adhere to the epithelial cells. van Cleef et al, showed up to 94% of those testing positive after working around livestock would test negative within 24 hours of the exposure [39]. As our study was cross-sectional, it was not possible for us to determine the duration of colonization for those participants with livestock exposure. We were able to assess how long it had been since the livestock workers last contact with livestock. For swine and cattle, it had been around 24–30 hours since last contact, so it is likely any *S. aureus* presence is true colonization, not contamination.

We also assessed the differences between livestock workers who were colonized and not colonized in the nares and oropharynx to assess any microbial differences that may influence whether a livestock worker was a carrier or not. While there were no significant differences in the nares, colonized livestock workers were significantly more likely to carry *Porphyomonas*, specifically *P. gingivalis* in the oropharynx while non-colonized livestock workers were more likely to carry *Atopobium* in the oropharynx. Studies have shown in the presence of *P. gingivalis*. *Streptococcus* spp. (which was significantly more abundant in colonized livestock workers) and other members of the oropharyngeal microbiome will aggregate with *S. aureus*. Meaning even if *S. aureus* is a contaminant instead of a true colonizing species in the oropharynx, when *P. gingivalis* is present, *S. aureus* may adhere to the biofilms created by the other microorganisms. *P. gingivalis* acts as a ‘coaggregation bridge’ mediating the attachment between two species that would not otherwise aggregate [40]. Additionally, the presence of *P. gingivalis* in the oral cavity is associated with poor oral health and hygiene [41]. In our study, livestock workers were significantly less likely to brush their teeth daily (*P* <0.001, data not shown) which has been linked to poor oral health and gingivitis [42]. However, we were not able to identify an association between oral hygiene and any differences in the microbiota in our study, likely do to small sample sizes and an inadequate measure of oral hygiene. While this observation requires a great deal of further study to better understand and quantify the relationship between *P. gingivalis* and *S. aureus* as our numbers are small, the improved oral hygiene may help reduce oropharyngeal *S. aureus* colonization.

This study was cross-sectional. As such, we were unable to assess any change in the microbiome and how those changes may impact *S. aureus* colonization. It’s also highly possible some of the intermittent *S. aureus* colonizers were mistakenly classified as not colonized [4] which could mask any differences in the microbiomes.

## Conclusions

The microbiome likely plays a crucial role in *S. aureus* colonization, and while we were unable to identify any differential species, we did observe a difference in the beta diversity between the colonized and non-colonized subjects. There is a great deal of difference between colonized livestock workers and non-livestock workers as well as differences between colonized and non-colonized livestock workers, specifically, *P. gingivalis*. Improving oral hygiene may help to reduce oropharyngeal carriage of *S. aureus* and MRSA in livestock workers.

Longitudinal studies on larger populations are needed to better characterize the relationship between the microbiome and *S. aureus* carriage. Understanding community compositions is a necessary start understanding colonization, but does not provide a full picture of the relationship between the microbiome and colonization with pathogenic organisms such as *S. aureus*. Studies assessing the potential interactions between the host, the microbiome, and *S. aureus* are needed in order to successfully manipulate the microbiome to inhibit *S. aureus* colonization.

### Funding

This publication was supported in part by Grant Number 5 U54 OH007548-11 from CDC-NIOSH. Its contents are solely the responsibility of the authors and do not necessarily represent the official views of the CDC, NIOSH, or the Great Plains Center for Agriculture Health. The funding organization had no influence on the study design, data analysis, interpretation of data, or writing of the manuscript.

## Acknowledgments

The authors would like to thank Dr. Patrick Breheny for his invaluable guidance as well as the University of Minnesota Genomics Center for conducting the sequencing.

